# A sequence motif for DNA double-strand break and telomere healing during programmed DNA elimination

**DOI:** 10.64898/2026.03.06.709695

**Authors:** Jansirani Srinivasan, Marro Agbaga, Vincent Terta, Brandon Estrem, James R. Simmons, Ryan Oldridge, Mehwish Iftikhar, Abigail West, Hannah Lam, Thomas C. Dockendorff, Jianbin Wang

## Abstract

Programmed DNA Elimination (PDE) is an exception to the paradigm of genome integrity, removing selected DNA sequences during development. PDE is observed in dozens of metazoan species from diverse phyla, but its molecular mechanisms and biological significance in most metazoa remain largely unknown. During PDE in the nematode *Oscheius tipulae*, DNA double-strand breaks (DSBs) are generated at subtelomeric regions, followed by the loss of DNA at chromosome ends and the healing of the DSBs by *de novo* telomere synthesis. DSBs occur at a 29-bp degenerate palindromic <u>S</u>equence <u>F</u>or <u>E</u>limination (SFE) motif. We determined the sequence requirement for DSB generation and demonstrated that the conserved GGC/GCC sites are used for neotelomere formation. Introducing the SFE into a retained DNA region adjacent to a native SFE induces DNA cleavage, telomere healing, and loss of additional DNA between the two SFEs. Moreover, insertion of the SFE in the middle of the sex chromosome splits it into two functional somatic chromosomes, demonstrating that the function of SFE is not necessarily constrained by its genomic location. Overall, our data show that the SFE motif is both necessary and sufficient for the generation of DSBs and healing of DSB ends via telomere addition, providing molecular insights into the mechanisms of metazoan PDE.

## Introduction

Genome integrity is paramount to the development and reproduction of an organism. However, certain species change their genome through programmed DNA Elimination (PDE) by eliminating a portion of their DNA in somatic cells during development (1–6). PDE occurs in various forms and has been identified in a growing number of organisms from diverse phyla (7–14). Organisms with PDE can be divided into two major groups based on how their DNA is lost. One group, including ciliates, nematodes, and copepods, uses DNA double-strand breaks (DSBs) to fragment the chromosomes, followed by repair of the DSBs and selective segregation of the retained DNA (7,15). The other group, encompassing (but not limited to) certain species of insects, fish, and birds, eliminates entire chromosome(s) without inducing DSBs (12,14).

For PDE that involves DSBs, several key questions on the mechanism(s) of PDE include: 1) how the break sites are identified; 2) how the DSBs are generated; and 3) how the DSBs are repaired to circumvent the DNA damage response, chromosome fusions, and genome instability. Single-cell ciliates use DSBs and constitute a well-established model and have led to major mechanistic insights into their PDE processes (16,17), including the domesticated transposons that generate the DNA breaks (18), small RNAs for marking retained or eliminated DNA (19,20), and telomerase for *de novo* telomere synthesis (15,21). In contrast, little is known about these processes in metazoa (1,2). A major model for metazoan PDE has been the parasitic nematode *Ascaris* and its relatives (ascarids) (1,2,7,22,23). In ascarids, DSBs are generated at the chromosomal breakage region (CBR) that spans 3-6 kb, and the broken DNA ends are healed through telomere addition that is independent of specific sequences (24,25). However, the molecular machineries responsible for CBR recognition and DSB generation in ascarids remain elusive (7,26,27).

Recently, PDE has been identified in several nematodes amenable to genetic manipulation, providing new opportunities for functional and mechanistic studies (28). One such species is the free-living *Oscheius tipulae* (29,30). *O. tipulae* eliminates only 0.6% of its genome that contains 112 genes, most of which are silenced in the germline or expressed at a low level in early embryos (29). The amount of the eliminated DNA and the number of genes are small compared to other known nematodes with PDE (7). *O. tipulae* has several features that make it a good model to carry out in-depth molecular and genetic analyses of metazoan PDE (29). It has a small genome [60 Mb, with complete genome assembly (30) and comprehensive transcriptomes (29)], reproduces mostly by self-fertilization (androdioecy), has a rapid generation time (3-4 days), a transparent body, and is easy to analyze microscopically (31). In addition, large quantities can be cultured for biochemical and molecular studies, and CRISPR tools have been adapted for this nematode (29).

We previously identified a sequence motif in *O. tipulae*, known as the Sequence For Elimination (SFE), at the junctions of the retained and eliminated DNA at all 12 chromosome ends (29). The SFE is a 29-bp degenerate palindromic motif that has highly conserved blocks as well as more variable regions. Independent disruption of three SFE sites using CRISPR led to the PDE failure at the altered SFEs, demonstrating an essential role for the SFE in PDE (29). Our molecular analysis of the SFE sites indicates that DSBs occur within the motif, and that the DSB ends are resected to generate a 3’-overhang necessary for *de novo* telomere addition (29). Together, our data suggest that the SFE is required for both DSB generation and neotelomere formation during *O. tipulae* PDE.

Here, to determine the key sequence elements of the SFE, we carried out genetic analyses by mutating the SFE sequences. These studies identified nucleotides within the motif that are essential for PDE, including specific sequences needed for telomere addition. To determine if the position of the SFE in the chromosomes is important to execute PDE, we inserted SFE sequence into various locations of the genome and observed that the SFE can direct DSBs and telomere addition in diverse locations in the genome. Together, our data suggest that the SFE is necessary and sufficient to trigger PDE. The function of SFE as a discrete genetic element for PDE may also influence the organization of the *O. tipulae* genome over evolutionary time.

## Results

### SFE consensus sequence can trigger PDE

*O. tipulae* eliminates 349 kb of non-telomeric DNA, along with all germline telomeres that cap the chromosome ends (29,30). We previously demonstrated that a 29-bp degenerate palindromic SFE sequence is needed for chromosome breaks and telomere addition during PDE (29). By aligning the 12 SFEs, we derived a consensus SFE sequence (Fig. 1A). We first evaluated whether a consensus sequence based on the SFEs could also induce PDE. Given the significant variations among these SFEs (Fig. 1A), testing the consensus is more practical than individually testing each SFE. Since the SFE is largely palindromic, we aligned the SFEs based on the positions of the most conserved GGC/GCC nucleotides (Fig. 1A). This alignment produced a palindromic motif with three or four less-conserved base pairs between the conserved GGC and GCC. Sequence analysis of 12 alternative SFEs in the eliminated regions (29) and 42 SFEs from several wild isolates showed they all have three base pairs between the GGC and GCC, indicating that 3-bp spacer is conserved. Thus, we used the 3-bp spacer consensus in this study (Fig. 1B).

**Figure 1.**
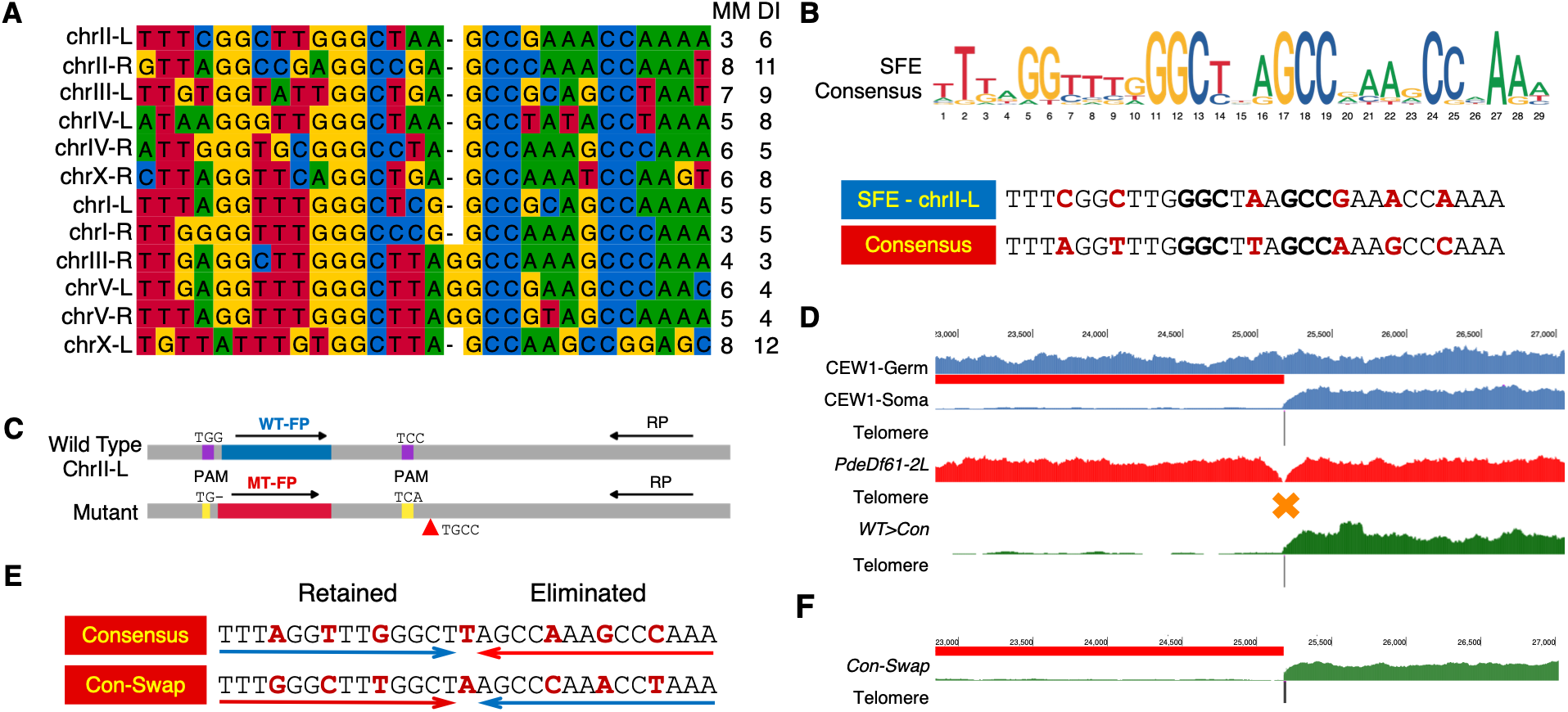
An SFE consensus sequence can trigger PDE. (**A**) Alignment of 12 sequences from *O. tipulae* at the DNA break sites in the wild-type CEW1 strain. The alignment was generated using Jalview (75) with manual refinement. A 1-bp gap was introduced in the middle of nine sequences to align with the other three (chrIII-R, chrV-L, and chrV-R) using the flanking ultra-conserved GGC/GCC sequences. The number of mismatch (MM) base pairs within each palindromic sequence is indicated at the right. The number of diverged (DI) base pairs compared to the consensus is also indicated. (**B**) A sequence motif derived from the SFE alignment in Figure 1A, after removing the unaligned middle G nucleotide in chrIII-R, chrV-L, and chrV-R. Below is a comparison of the chrII-L SFE and the consensus sequence. The conserved GGC/GCC sequences are in bold, and the divergent nucleotides are highlighted in red. (**C**) Schematic of the CRISPR genome editing strategy for replacing a wild-type SFE in ChrII-L with the SFE consensus sequence. Blue indicates the wild-type SFE, while red denotes the consensus SFE. Purple marks the PAM sites used for Cas9 cleavage, and yellow indicates mutated PAM sequences to prevent re-cutting by Cas9. The black arrow denotes the primers used for genotyping: the forward primers (FPs) are unique in wild type and mutant, while the reverse primer (RP) is common. The red triangle indicates an alternative design wherein we eliminated four nucleotides (TGCC; see Table S1) to ease detection of mutants by PCR. (**D**) Detection of PDE in mutants by genome sequencing. Read coverage is shown for wild-type germline and somatic cells (blue; top tracks), a mutant that removed 61 nucleotides including the SFE (red; *PdeDf61-2L*), and a mutant with the consensus SFE (green; *Pdesb29(WT-Con)*). Red bar marks the eliminated DNA. Telomere addition sites are shown in black. (**E**) Swapping of the left and right sides of the palindromic consensus sequence. The retained and eliminated sides of the SFE sequence in the consensus mutant (top) are underlined in blue and red arrows, respectively. The sharp ends of the arrows point to the break site. In the *Con-Swap* mutant (bottom), the retained and eliminated sides are swapped and marked with red and blue arrows, respectively. Changed bases are highlighted in bold and red. (**F**) The *Con-Swap* mutant triggers PDE. Genome sequencing of the *Con-Swap* mutant. Same legend as in Figure 1D.

We first replaced the wild-type SFE at the left end of chromosome II (chrII-L) with the consensus SFE using CRISPR (Fig. 1C). ChrII-L was chosen since only ~10 kb of non-telomeric sequence is lost at this end, and a mutant that failed to eliminate this 10 kb DNA did not produce any noticeable phenotype (29). Compared to the wild type chrII-L SFE, the consensus sequence differs by six nucleotides within the 29-bp motif (Fig. 1C). This difference is sufficient to allow the detection of the edited genome through PCR without changing any other sequences in the genome, except for the PAM sites to prevent CRISPR re-editing (Fig. 1C). Genome sequencing of a homozygous mutant revealed a wild-type pattern of PDE at the modified locus (Fig. 1D), indicating that the consensus sequence induced PDE, illustrating the sequence has the features required for SFE-mediated PDE.

While most nucleotides within the SFE are palindromic, the retained (left) and the eliminated (right) sides of the SFE show sequence variations in each individual SFE (Fig. 1A). For the SFE consensus, three out of the fourteen nucleotides on each side are not complementary (Fig. 1E). Notably, these three pairs of nucleotides that defy the palindrome complementarity are overall conserved in the 24 SFEs across the genome, suggesting the orientation of the SFE may play a potential role (29). To further investigate this, we swapped the two sides without altering their flanking sequences (Con-Swap), resulting in a 7-bp difference in the mutant (Fig. 1E). Genome sequencing analysis of the mutant with the swapped SFE revealed that it can still trigger PDE (Fig. 1F). This suggests that the two sides of the SFE consensus are interchangeable.

### Identifying regions of the SFE critical for PDE

To identify key sequence elements within the SFE, we mutated various regions of the consensus sequence focusing on the retained side since the two sides of the SFE are interchangeable (Fig. 1E-F). We adopted a targeted mutagenesis strategy in which we scrambled five 4-bp regions covering the first 16 bps of the consensus sequence, including the 3-bp spacer (Fig. 2A). We selected the four-base regions to ensure sufficient changes within each region while maintaining a manageable number of experiments. Although there are many possible permutations, each modified SFE sequence was designed to maximize the difference between the original and the scrambled sequences. Despite our concerted efforts, we were unsuccessful in creating an *O. tipulae* CRISPR mutant that alters the 1-4 nucleotides from TTTA to TATT (see methods for details). Nevertheless, we generated the other four mutants that cover most of the retained side of SFE (Fig. 2A). Genome sequencing of the homozygous mutants indicates mutated sequences within the more conserved regions (nucleotides 4-7, 11-14, and 13-16) of the SFE impact the ability to execute PDE, while changes to the less conserved region (nucleotides 7-10) did not impact PDE (Fig. 2B and Fig. S2).

**Figure 2.**
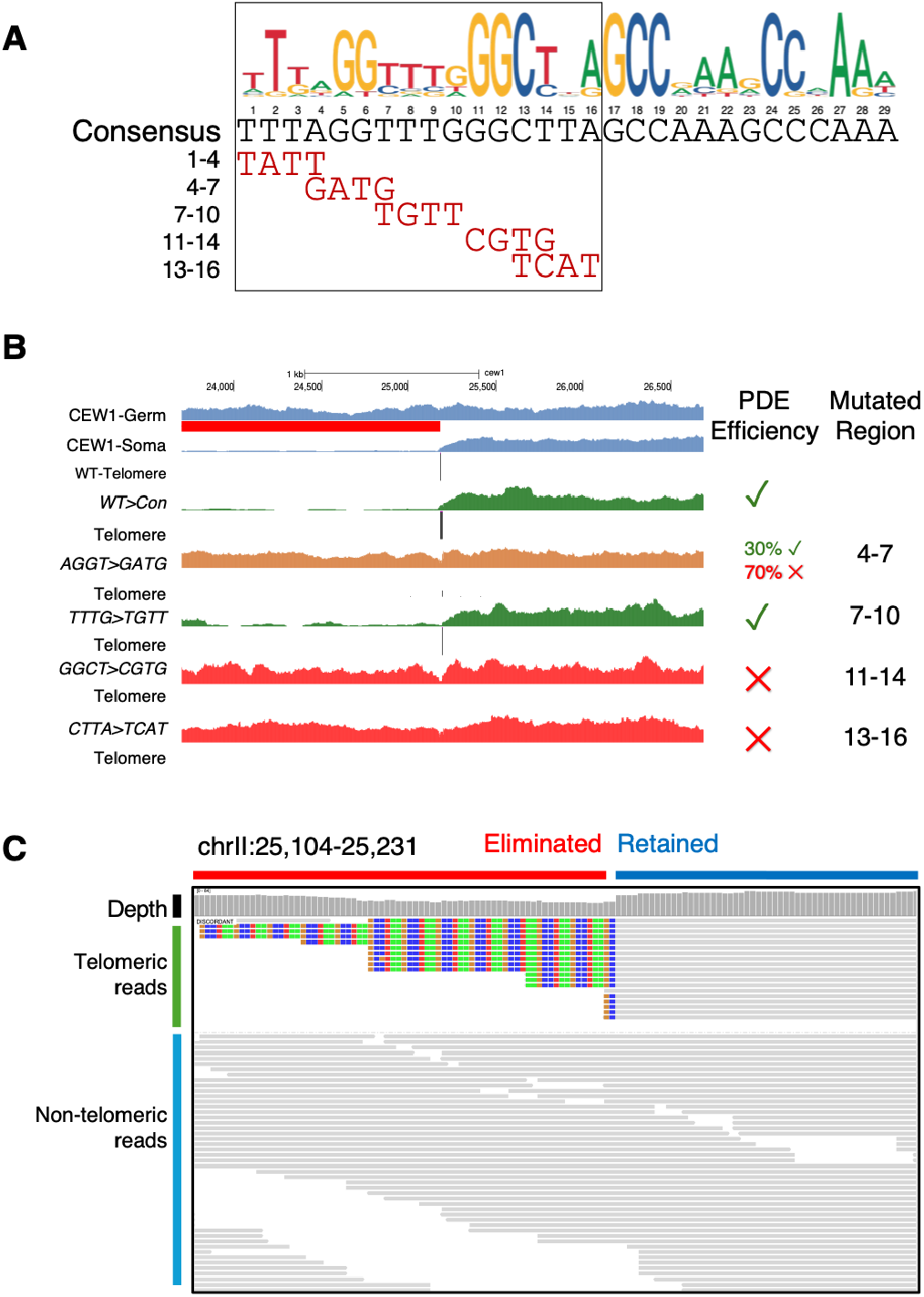
Functional analysis of the sequences within SFE required for PDE. (**A**) Scrambled 4-bp sequences covering the retained side of the SFE motif. Only the mutated nucleotides are shown. These mutated consensus sequences were used to replace the wild-type chrII-L sequence using the same strategy as shown in Figure 1C. (**B**) Genome sequencing revealed the contribution of the mutations to PDE. Same legend as in Figure 1D. Mutants that fail to undergo PDE are displayed in red, while those that successfully undergo PDE are shown in green. Mutants that partially undergo PDE (nucleotides 4-7, AGGT>GATG) are indicated in brown. The efficiency of PDE for each mutant, measured by the percentage of somatic reads with telomere addition, is indicated, with green ✓ indicating 100% efficiency and red ✗ indicating 0% efficiency. The changed regions are indicated at the right. Note that the mutant for nucleotides 1-4 is not available, despite multiple attempts to create one (see Methods). (**C**) Genomic reads in the partially functional AGGT>GATG mutant are shown in an IGV view. The horizontal red bar at the top marks the eliminated region, while the blue bar indicates the retained region. Reads with new telomere addition (TTAGGCs) are highlighted with colored bases and marked with a green vertical line at the left. Non-telomeric (unchanged germline) reads are marked with a blue vertical line at the left. See Figure S2E for the counts and percentages of germline reads for all 12 SFE sites.

Interestingly, while the AGGT>GATG mutant (nucleotides 4-7) exhibits impacts on DNA elimination, 29% (19/66) of the reads matching the mutated SFE had new telomeric sequences (Fig. 2C). In comparison, the other 11 wild-type SFE sites had ~95% of the reads with telomeres (the other ~5% are presumably germline reads that do not undergo PDE; see Fig. S2B). Thus, the AGGT>GATG mutation can still lead to PDE albeit at only ~30% efficiency (29%/95%, when factoring in the amount of germline reads in the sample). In contrast, no telomeric addition was observed in the other two mutants (nucleotides 11-14 and 13-16), suggesting a complete block of PDE. While we did not test all permutations of possible sequence mutations, this AGGT>GATG mutant suggests that some SFE-like sequences may exhibit a partial PDE phenotype, implying variations in the strength of the interactions between the motif and the presumed SFE-binding protein(s). Overall, our findings highlight the functional significance of conserved sequences within the SFE.

### The conserved GGC/GCC sites are used for neotelomere healing

The DSBs that initiate PDE need to be promptly healed by telomere addition to prevent other unintended DNA repairs or fusions (32). We previously showed that telomere addition occurs on the resected DSBs without bias on both the retained and eliminated sides of the break, immediately adjacent to the conserved GGC/GCC sequence (29). Since the resected DSBs have the GGC/GCC sequence at the ends of the 3’-overhang that are complementary to the telomeric repeat TTA<u>GGC</u>/<u>GCC</u>TAA (Fig. 3A), we hypothesize that the GGC/GCC could be used by telomerase RNA priming for new telomere synthesis (33). To test this, we mutated the consensus to alter the telomere-proximal GCC sequence on the eliminated side (nucleotides 17-19; Fig. 3A). If the mutated sequences can trigger DSBs, telomere addition might still heal the unchanged retained side of the break. If functional for DSB, the mutant would have minimal overall functional consequence since the DNA from the eliminated side would likely still undergo degradation. However, the mutants might allow us to assess the sequence requirements for telomere addition on the eliminated end of the break.

**Figure 3.**
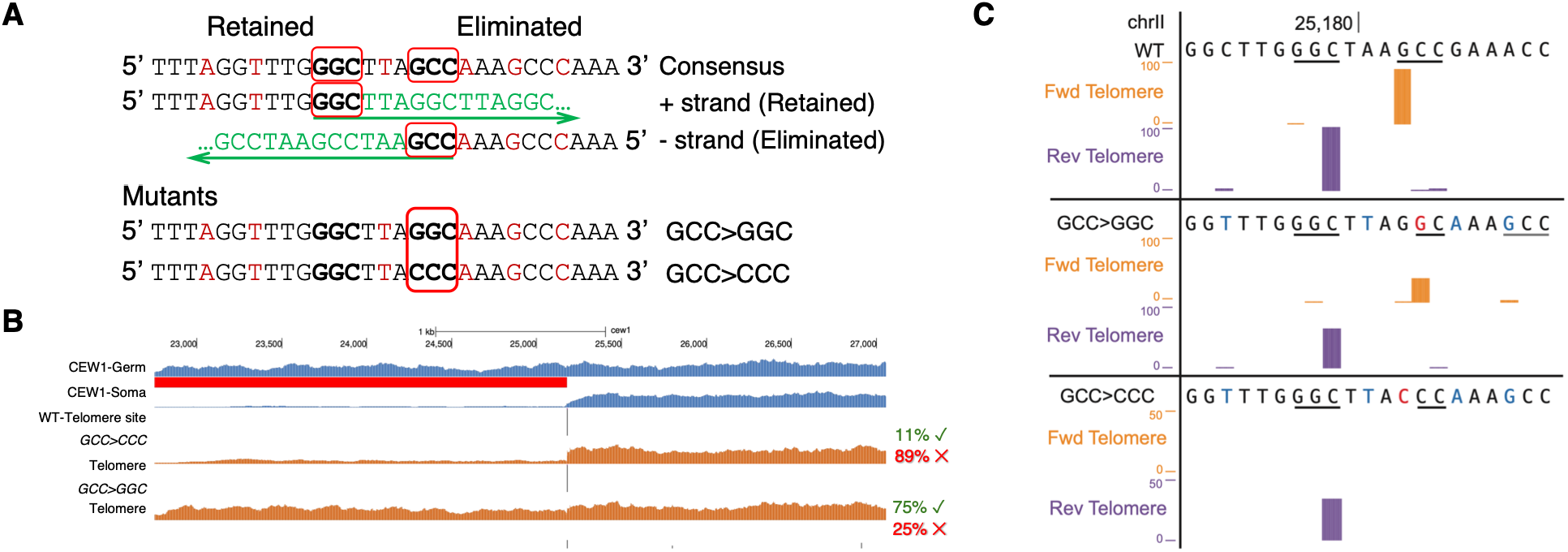
The conserved GGC/GCC sequences are the sites for neotelomere formation. (**A**) Telomere addition occurs after a DSB. During PDE, the germline SFE (consensus) is broken in the middle. The broken ends are healed by the synthesis of new telomeres at both the retained and eliminated sides. Telomere repeats (TTAGGCs) are colored in green, and the direction of the green arrows underneath the telomeric sequences point to the direction of telomere synthesis. The conserved GGC/GCC are highlighted in red boxes (top). Changes to the conserved GCC on the eliminated side of the SFE consensus in the mutants are highlighted in the red box (bottom). (**B**) Genome sequencing of the mutants. In the GCC to CCC mutant, the majority (~89%) of the reads at the SFE sites are germline, indicating the mutant largely prevented PDE. The rest ~11% of the reads have newly added telomeres (see Figure S3A). In contrast, in the GCC>GGC mutant, PDE proceeds largely as normal, although a higher (~29%) number of reads are non-telomeric reads (see also Figure S3B). Same legend as in Figure 1D. (**C**) New telomere addition sites in the wild type and mutants. DSB ends healed with telomere addition were enriched with END-seq in the wild type and the GGC mutant. Reads that have both telomeres and unique sequences were selected and mapped to the genome to determine the telomere addition sites; they were colored in orange for the forward strand and purple for the reverse strand. The upper panel shows the telomere priming sites in wild type using the same sequence (GCC in the forward strand and GGC in the reverse strand) at both sides of the DSB. The middle panel displays telomere addition in the GGC mutant, which also occurs at both sides. A two-nucleotide GC (in the forward strand) that matches the telomeric sequence is used on the eliminated side. Note that the downstream GCC site (in the forward strand) located six nucleotides away was not the preferred site for telomere addition. The bottom panel shows the telomere addition sites from whole genome sequencing of the CCC mutant. Due to the low level of PDE efficiency (11%) in the CCC mutant and the low amount of eliminated DNA in the mixed worms, we did not identify any telomere reads at the eliminated side. Note that the telomere reads are raw counts, so the scales are different.

Due to the likely importance of the GCC at nucleotides 17-19 (see Fig. 2), we chose to introduce a single nucleotide change to the GCC and created two such mutants (Fig. 3A). In the GCC>CCC mutant, sequencing of genomic DNA from a mixed worm population showed a small number of reads (4/41 or 10%; see Fig. S3A) with telomere addition at the mutated chrII-L site. In comparison, an average of ~91% of reads at the other 11 SFE sites have new telomere addition (see Fig. S3B). This suggests that the GCC>CCC mutant largely blocked PDE (the PDE efficiency is ~11% [10%/91%]). In contrast, genome sequencing showed that the GCC>GGC mutant has 43/64 (67%; see Fig. S3C) of the reads with new telomere at the chrII-L site, while the other 11 sites have ~90% of telomeric reads (see Fig. S3D), indicating that the GCC>GGC mutant largely retained the ability to trigger PDE (the PDE efficiency is ~75% [67%/90%]). Thus, the two single-base mutants with altered GCC sequence can still trigger DSBs and heal the retained sides with telomere addition at varying efficiencies.

We next focused on the GCC>GGC mutant that has a higher PDE occurrence (75%) to determine if and at which position(s) new telomeres are added to the eliminated side of the DSBs. To enrich for the to-be-eliminated DNA, we used staged early embryos (~8-32 cells), since PDE occurs at 8-16 cell stage(29) and the DNA to be eliminated is not fully degraded for a few cell cycles post PDE. To further enrich the signal, we also applied END-seq, which detects DSB ends that include newly added telomere ends (29,32). Our END-seq results show that the majority (30/33 or 90%) of the telomere addition sites at the eliminated side are shifted by one nucleotide compared to the wild type (Fig. 3C). This new site contains a GC dinucleotide that matches the telomeric sequence (<u>GC</u>CTAA). The other three observed telomere addition sites were at much lower frequencies, including a downstream GCC sequence six nucleotides away that matches the telomere <u>GCC</u>TAA (Fig. 3C). Overall, these data confirm that the conserved GGC/GCC sites are the main sites for telomere priming. Furthermore, they suggest that minimal trimming occurred at the 3’ end of the DSBs before telomere addition, and that the telomere addition machinery favors sites closer to the DSBs over sites with a perfect 3-nt (GGC/GCC) priming sequence (see discussion).

### The SFE consensus can function at different genomic locations

All the mutants described above were created by altering the endogenous sequence at the chrII-L site. To determine if the SFE function is also dependent on its native genomic location, we chose to insert the SFE consensus towards the retained side of chrII-L. Since a functional SFE in the retained region could eliminate additional sequences potentially essential for somatic function and thus lead to lethality, we first screened for prospective insertion sites by analyzing RNA-seq data (29) to avoid eliminating genes that are highly expressed in somatic tissues (Fig. 4). Due to the compact genome of *O. tipulae* (29,30), regions available for insertion of the SFE without potentially eliminating somatic genes are limited. Nonetheless, candidate insertion sites were selected on the retained side of the chrII-L SFE at about 1 kb and 10 kb distal to the native site (Fig. 4). Using CRISPR, we created independent homozygous mutants harboring the SFE consensus for both sites.

**Figure 4.**
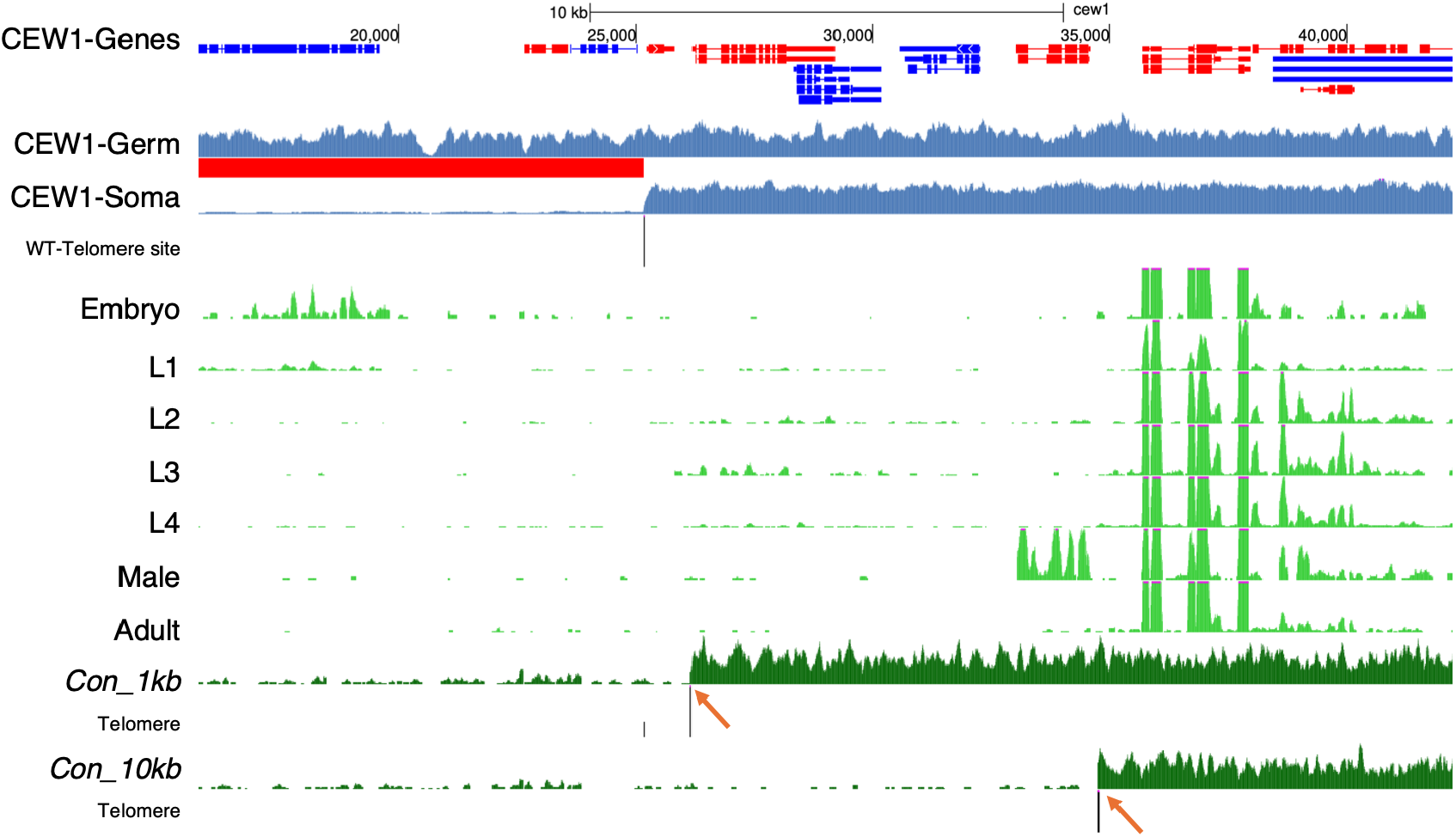
The SFE consensus can function at different genomic locations. Genome sequencing of two mutants with an insertion of the SFE consensus at 1 kb or 10 kb into the retained region of chrII-L. Same legend as in Figure 1D. Gene models at the top show genes transcribed in the forward strand colored in red and in the reverse strand in blue. Normalized RNA-seq data from embryos, larvae (L1 to L4), and adults (hermaphrodites and males) are shown in light green tracks to indicate that no actively transcribed genes in the somatic cells and one active gene in the male are located in either the eliminated regions or within the 10 kb retained region adjacent to the canonical SFE. Red arrows indicate the inserted SFE sites at 1 kb and 10 kb, where new telomeres were added after the SFE triggered DSBs.

Insertion of SFE consensus at both the 1-kb and 10-kb sites leads to PDE and the loss of additional DNA (Fig. 4). We also observed in the 1-kb mutant, the native SFE site still undergoes PDE (Fig. 4 and Fig. S4), suggesting a functional SFE does not appear to inhibit the function of a neighboring SFE 1 kb apart. We believe the lack of telomere signal at the canonical chrII-L site in the 10-kb mutant (Fig. 4) is likely due to the lower sequencing depth and the degradation of the eliminated DNA, as seen in the other alternative SFE sites where little telomere addition is detected (data not shown). Furthermore, the 29-bp SFE consensus was flanked by as little as 4 base pairs of original flanking sequence (Table S1). The data reinforce the idea that the 29-bp SFE functions as an independent unit for DNA break and telomere addition during PDE.

### The SFE is sufficient for PDE

Although the 1-kb and 10-kb sites are independent of the native chrII-L site, they are close to each other, and like the native site, they are all located at the end of the same chromosome. Therefore, a mechanism not relying on specific sequences, such as 3D genome organization (34–36), or a location near the telomeres could be required for PDE. We next inserted the SFE consensus in the middle of a chromosome. As most nematodes have holocentric chromosomes (37,38), a functional SFE in the middle of a chromosome would split the chromosome and lead to karyotype changes but would be otherwise viable, as commonly observed in other nematodes with PDE (7,39). A typical feature of a nematode chromosome is the presence of euchromatin in the middle and heterochromatin at the ends (37,40). To facilitate the selection of a site that may lead to two somatic chromosomes with heterochromatic ends, we first obtained histone modification data (CUT&RUN) for H3K9me3, a heterochromatin mark, in *O. tipulae* embryos (Fig. 5A). We found an 80-kb window flanked by two major peaks of H3K9me3 in the middle of the sex chromosomes (chrX:5,900,000-5,980,000), providing an ideal site for a potential insertion of the SFE (Fig. 5A).

**Figure 5.**
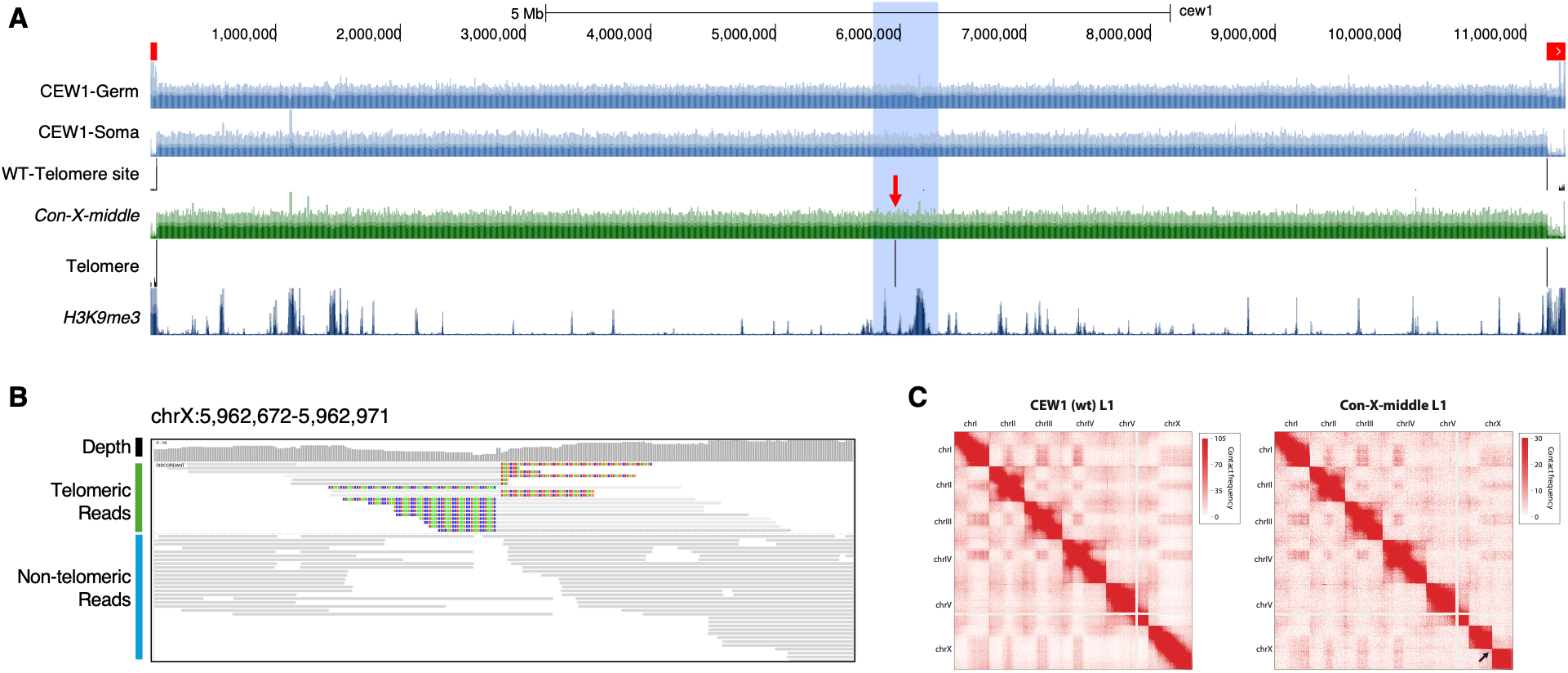
Insertion of SFE in germline sex chromosome leads to two somatic chromosomes. (**A**) Genome browser view of the sex chromosome with the SFE consensus inserted into the middle of the chromosome (blue highlighted region; red arrow indicates the insertion site). Note the insertion site is in between two major H3K9me3 peaks. The genome sequencing of the mutant, *Con-X-middle*, is shown in green. Note the detection of telomere addition at the SFE insertion site. Same legend as in Figure 1D. (**B**) Genomic reads in the *Con-X-middle* mutant were shown in an IGV view. Same legend as in Figure 2C. Both sides of the DSB induced by the inserted SFE are healed with new telomere addition. As expected, telomere sequences on the forward or reverse strand appear with alternative color patterns. (**C**) Hi-C data show that the *Con-X-middle* mutant larvae L1 has seven somatic chromosomes, one more than wild type. The black arrow indicates the position of the newly inserted SFE in the middle of the sex chromosome.

We created a homozygous mutant that harbors the SFE consensus in the middle of the X chromosome. Genome sequencing of a mixed worm population from this mutant revealed no loss of DNA at the inserted site (Fig. 5A). However, telomere sequences (18 out of 21 reads, ~86%) were found at both sides of the DSB generated in the middle of the SFE (Fig. 5B). The remaining ~14% of reads are likely germline reads that do not undergo PDE. This telomere addition ratio is consistent with the ratio at the 12 native chromosome ends (111 out of 138 total reads; ~80%), suggesting that the introduction of an SFE in the middle of the X chromosome induced PDE at 100% efficiency. Overall, the data indicate that the SFE consensus is sufficient to induce PDE when introduced to a different genomic location.

To further confirm that the X chromosome has been split into two somatic chromosomes, we performed Hi-C in a somatic sample (larvae L1, over 99% are somatic cells). The Hi-C map showed that the mutant L1 contains seven chromosomes (Fig. 5C). Notably, the Hi-C block corresponding to the wild-type X chromosome is split into two blocks in the mutant. These data indicate that introducing SFE in the middle of the chromosome splits the sex chromosome and alters the somatic karyotype. Notably, the mutant is viable, fertile, and exhibits no apparent phenotypes. Together, our data suggest that the SFE is necessary and sufficient for generating DNA breaks and neotelomere formation during PDE.

### Sequence-independent factors that may contribute to PDE

Since the SFE motif exhibits high sequence variation (Fig. 1A), we speculate that there might be other sequences in the genome that may closely resemble the SFE. We manually examined the top 137 FIMO-predicted hits (score cutoff >= 60) (29) that possess the characteristic features of a conserved SFE: containing the conserved GGC/GCC with a 3 to 4 bp spacer in between. Four potential sequences were identified as plausible SFEs because they share these features (Fig. 6A), and some appear to be similar in sequence divergence compared to the SFE consensus than other known functional SFE sites (such as chrX-L). However, we did not detect any signs of DNA breaks or telomere addition at these four FIMO sites. These data suggest either that these sites are too diverged and lack some uncharacterized sequence feature of the SFE or that they have other sequence-independent features that prevent them from executing PDE.

**Figure 6.**
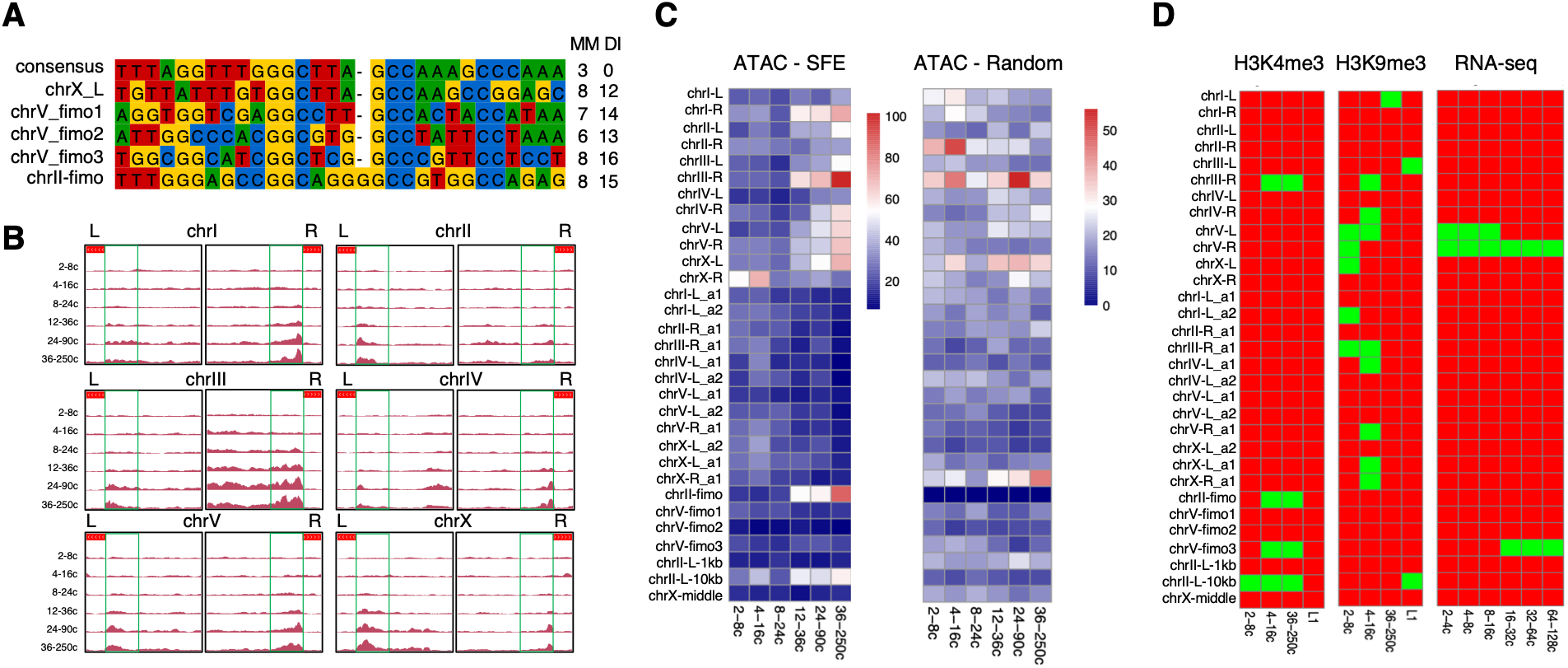
Non-sequence features associated with SFE sites. (**A**) Alignment of four sequences predicted by FIMO with high similarity (cutoff score: 60) to the consensus SFE. The consensus SFE and the chrX-L SFE, which has eight mismatches in the palindromic pairs, were included for comparison. Same legend as Figure 1A. (**B**) ATAC-seq profiles for all 12 SFEs from developmental stages before, during, and after PDE. For each SFE site, we included 500 bp of the eliminated region (marked with a red bar at the top) and 2,500 bp of the retained region. “L” denotes the left and “R” the right end of the chromosomes. Green boxes mark regions that show consistent increased chromatin accessibility after PDE. (**C**) Heatmap of ATAC-seq signal at the SFE sites (flanking 500 bp at each side). The canonical SFE sites (top 12) show a consistent increase in accessibility after PDE, while the 12 alternative SFEs (middle 12) and the FIMO predicted sites show a decrease of signals due to the loss of the DNA over time. Note that the alternative SFEs display no accessibility after PDE due to loss of sequence in the somatic cells. Randomly selected 1-kb genomic regions (ATAC-control, identified by BEDTools; see methods) were used for comparison. Heatmap scale: red = high accessibility, blue = no accessibility. (**D**) Histone modifications and RNA expression data at the SFE sites were shown as signal (green, defined as a peak region) and noise (red, defined as a non-peak region). See methods for peak calling. Data for the same genomic regions and the same order as in Figure 6C (1 kb surrounding the SFE-related sites) were shown.

To discern if other features could be associated with the execution of PDE, we compiled 27 known sites that can trigger PDE, including the 12 canonical SFEs, 12 alternative SFEs, chrII-L 1-kb, chrII-L 10-kb, and chrX-middle, as well as the mutated sequences that can still trigger PDE (Fig. 2 and Fig. 3), and compared them to the four FIMO sites that did not cause PDE. Since the palindrome is a main feature of the SFE, we examined the number of mismatches in the palindromic pairs to determine if the sequence complementarity is associated with the function of SFE. This analysis showed significant variations ranging from 3 to 8 mismatches across the functional SFEs (Fig. 1A). No significant differences were found on the number of mismatches between the SFEs that trigger PDE (average 5.5 mismatches) and the four FIMO predicted sequences that do not (ranging from 6 to 8 mismatches, with an average of 7 mismatches; see Fig. 1A and Fig. 6A). These results suggest that the sequence complementarity between the two sides of the palindrome is unlikely required for the PDE function (see discussion).

We next evaluated the chromatin accessibility (ATAC-seq), selected histone marks (CUT&RUN on H3K4me3 and H3K9me3), and RNA expression level (RNA-seq) at the SFE and FIMO sites. We found little correlation between these features and the SFE sites (Fig. 6B-D). For example, ATAC-seq showed that although the 12 native sites became more accessible after PDE, there is little difference in the chromatin accessibility between the functional SFE and the non-functional FIMO sites before and during PDE. Also, no pattern emerged in the active histone mark H3K4me3, the repressive mark H3K9me3, or the RNA-seq data that can distinguish the 27 sites that trigger PDE from the four FIMO sites that do not (Fig. 6B-D). Thus, although only a limited number of sequence-independent datasets were evaluated, none of them are linked to SFEs.

## Discussion

DNA double-strand breaks (DSBs) are generated in many species to initiate programmed DNA elimination (PDE) (1,2,6). The repair of these DSBs is crucial to genome stability. Key questions include how the DNA break sites are identified, how the DSBs are generate, and how the DSBs are recognized by specific repair pathways (7,15,41). To address these questions, we used a newly established genetic model of metazoan PDE, *Oscheius tipulae*, to characterize the SFE motif and its sequence requirements for DSB and telomere healing. We found that the consensus sequence of the 29-bp SFE motif in *O. tipulae* is sufficient to promote both the formation of the DNA break and its subsequent repair through neotelomere synthesis. Insertion of this motif at various genomic locations can trigger DSBs and telomere healing, including the partitioning of the sex chromosome into two functional somatic chromosomes.

### Sequence features of a functional SFE

Our genomic analyses reveal substantial variations among the functional SFE sequences. We generated a consensus motif and demonstrated that it was sufficient to induce DSBs and neotelomere formation during PDE. We then carried out directed mutational and functional analyses of the consensus motif. We showed that mutations within a less conserved region (nucleotides 7-10) showed no disruption on DSBs, neotelomere formation, and DNA loss (Fig. 2). For nucleotides 4-7 that contain a more conserved GG sequence, the mutation remains partially (30%) functional (Fig. 2B). Even for one of the most conserved sequences, nucleotides 17-19 (GCC), that are believed to be responsible for priming during the initiation of new telomere synthesis, a single nucleotide change can still partially (11% for the GCC>CCC mutant and 75% for the GCC>GGC mutant) retain the functionality of the SFE (Fig. 3). Furthermore, a reciprocal exchange of the left and right side of the consensus SFE can also trigger PDE (Fig. 1E-F). These findings, together with the known functionality of alternative SFEs and SFEs from various wild isolates (29), suggest a high degree of tolerance for variation within this 29-bp motif for PDE.

Meanwhile, mutations towards the center of the motif (nucleotides 11-14, 14-16, and 17-19) have a more pronounced impact on the outcome of PDE (Figs. 2 and 3). Given that the DSB occurs at the center of SFE, and the adjacent GGC/GCC is utilized for telomere priming (Fig. 3), it is plausible that the conservation of the GGC/GCC sequence is reinforced by the necessity to effectively heal the broken ends. The conservation of the SFE sequence may have evolved with additional sequences (such as the less conserved GG at nucleotides 5-6) that could serve as backup sites for telomere addition. Indeed, we observed that in the wild type, although at a much lower frequency (~2%), dinucleotides within the SFE that match the telomeric sequences are also used as new telomere addition sites (29). The constraint that requires telomere priming sites to be within the SFE sequence could potentially enhance and eventually fix the use of the motif in the population. Likewise, proteins required for DNA cleavage may have co-evolved with the SFE motif. Thus, we believe that the conservation of the SFE sequences may be largely shaped by the sequence requirement needed for healing of DSBs using telomere addition machinery. Consistently, several motif sequences identified in other nematodes with PDE also appear to contain the conserved GGC/GCC sequences (39), highlighting the crucial role of telomere addition in nematode PDE.

We demonstrated for the GCC>GGC mutant, that rather than trimming the double-strand DNA (dsDNA) and searching for a perfect site for telomere priming, the telomere-healing machinery prefers the closest suitable site that has a complementary dinucleotide (GC) to the presumptive telomerase RNA (Fig. 3). This suggests that minimal or no trimming of dsDNA is needed for telomere formation following a DNA break. This is consistent with our previous findings that the broken ends immediately undergo single-strand resection to generate a 3’-overhang structure for telomere addition (29). These findings underscore the significance of the position of a DSB over the exact priming sequences for telomere addition during *Oscheius* PDE. This could be attributed to the immediate necessity of repairing the DSB with telomere addition before other repair pathways can engage with the broken ends, potentially leading to fusion, recombination, and genome instability. Alternatively, machineries for end trimming or other repair pathways may not be active or available at the sites of SFE-induced DSBs during PDE. Consistent with this, exogenous DSBs introduced using X-rays did not appear to trigger their healing through telomere addition in another nematode, *Ascaris*, that undergoes PDE (32). It would be interesting to investigate how the choice of DNA repair pathways is determined and regulated for DSBs associated with PDE vs. non-PDE DSBs.

The length of the SFE motif and flexibility of the sequence within the motif are reminiscent of homing endonuclease recognition sequences, which typically range from 20 to 30 bp with extensive variations (42,43). The interactions between homing endonucleases and their targets can tolerate some sequence degeneracy within their recognition sequence, although this can impact cleavage activity (44). This is similar to some of the SFE mutant sites we observed (nucleotides 4-7 and 17-19) that have reduced frequencies of DSBs and telomere addition (Figs. 2 and 3). However, unlike the palindromic features of the SFE (albeit degenerate), homing endonuclease recognition sites are often more asymmetric (45). It’s tempting to speculate that the presumed protein that recognizes the SFE could, on its own, cut the DNA, similar to homing endonucleases. Alternatively, other DNA-binding protein(s), such as zinc finger proteins, that may or may not have nuclease activity, could also recognize the SFE. They could generate DSBs themselves or recruit an endonuclease that can cut at the middle of the SFE sequence. It’s also possible that a transposase capable of recognizing relaxed DNA substrates, like the one recently indicated in the ciliate *Oxytricha* PDE (46), might be used in *O. tipulae* PDE.

The SFE is likely recognized by a trans-acting factor. Since the consensus sequence can trigger PDE (Fig. 1D) but is absent in the wild-type genome, it suggests that the SFE function doesn’t rely on precise complementary nucleotide sequences such as small RNA targeting. Instead, it implies that a DNA-binding protein recognizes the SFE through a limited set of sequences within it. Swapping the left and right sides of the SFE didn’t affect its PDE function (Fig. 1F), suggesting that the presumed DNA-binding protein might interact with the entire 29-bp SFE sequence. Alternatively, two DNA-binding proteins, each interacting with one side of the SFE sequences similarly and independently, could dimerize to perform their PDE function. Furthermore, the significant variations in the mismatches within the palindromic sequence (two functional SFE have 8 mismatches out of a total of 14 nucleotides; Fig. 1A) suggests it’s unlikely that the PDE function relies on the formation of secondary structure through sequence complementarity between the two sides of the palindromic sequences. Instead, this observation aligns with the idea that the DNA-binding protein(s) can interact with the two sides separately, likely through the formation of a protein dimer.

### Functional and evolutionary significance of the SFE

Since the SFE is both necessary and sufficient for DSB formation and telomere healing, its presence in undesired regions could pose a significant threat to the genome. For instance, an SFE further into the retained regions could eliminate genes that are essential for somatic function. Moreover, splitting a chromosome at an inappropriate location could result in a harmful karyotype that could jeopardize the species’ survival. Interestingly, sequence analysis of the wild type revealed that SFEs and SFE-like sequences (predicted by FIMO) are highly concentrated at the ends of the chromosomes (29). Thus, their absence in internal chromosome regions may have been selected, and this distribution serves as a fail-safe mechanism, ensuring the elimination of the chromosome ends while not triggering DSB formation at other genomic regions. This highlights the pivotal role of the SFE and PDE in shaping the sequence organization of the genome through evolution, suggesting a functional significance of PDE.

With comprehensive chromosome end remodeling in *O. tipulae* PDE, genes essential for somatic functions must avoid the ends of the chromosomes. This naturally demarcates the genome into two regions: the retained regions where essential somatic genes must reside, and the eliminated regions where germline-only or even soma-toxic genes can occupy. The creation of these two genomic compartments could provide several benefits. First, the reduced constraints of the eliminated regions suggest that they could serve as a reservoir for novel genes. This could be advantageous for organisms, providing an additional layer of evolutionary possibility to generate variants (47–50). Second, the machinery for the SFE function ensures that SFE sequences are not placed within the retained regions that could lead to lethality by eliminating essential genes in somatic cells. Thus, the SFE may be considered as a boundary element that limits essential somatic genes from spreading into the eliminated regions. Consequently, the *O. tipulae* SFE system effectively constrains itself to the ends of the chromosomes, safeguarding the retained genome from being altered by the SFE system.

### Comparisons to other species that use DSBs to execute PDE

Many other nematodes undergo PDE (7). In parasitic ascarids (22,51), including *Ascaris, Parascaris, Baylisascaris*, and *Toxocara*, telomere addition is sequence-independent and constrained within a large 3-6 kb regions known as chromosomal breakage regions (CBRs) (24,25). Notably, *Ascaris* CBRs lack a conserved motif or other sequence features, nor are they associated with small RNAs, R-loops, or histone marks (27,52,53). However, *Ascaris* CBRs are more chromatin accessible during PDE, suggesting alternative mechanisms are used to recognize and execute PDE (24–26). In contrast, free-living nematodes with PDE, including other *Oscheius, Auanema*, and some *Caenorhabditis*, a motif similar to the SFE at or near their break sites (39) is present, suggesting they may use a similar mechanism. Overall, these differences highlight the diversity in DSB sites used for nematode PDE: some rely on species-specific sequences, whereas others use a sequence-independent mechanism.

In ciliates, both sequence-dependent and sequence-independent mechanisms are used for DSBs during PDE. For instance, in *Tetrahymena thermophila* and closely related species, a 15-bp consensus chromosome breakage sequence (CBS) is present in the germline genome at least 200 times (54). The CBS is both necessary and sufficient for directing chromosome fragmentation (55). Interestingly, the CBS sequence in *T. thermophila* also includes a conserved hexanucleotide (CCAACC) that matches its tandem telomere sequence (CCCCAA). However, there’s a significant difference between CBS in *T. thermophila* and SFE in *O. tipulae*. In *T. thermophila*, trimming occurs to DSB ends at the CBS to various lengths (4-30 bp) before telomere addition occurs (56). In contrast, telomere addition occurs immediately at the DSB end in *O. tipulae* (Fig. 3). Furthermore, CBSs are highly conserved, with 50% of them matching the exact 15-bp consensus sequence, 40% having a single base difference, and 10% having two base variations (54). In contrast, the *O. tipulae* SFEs are highly divergent, with 3-12 bp (average of 7 bp) deviating from the 29-bp SFE consensus (Fig. 1A).

Interestingly, a recent study showed the *Tetrahymena* CBS are found in most of the subtelomeric regions that lead to DSBs and the loss of germline telomeres (57), similar to end remodeling in nematode PDE (24,29,39). Likewise, the *Euplotes* ciliates possesses a 5-bp motif sequence in the vicinity of most fragmentation and telomere addition sites in the somatic genome (58,59). In contrast, no consensus sequences adjacent to chromosome fragmentation sites were found in *Oxytricha* (60) and *Stylonychia* (61), although a recent study in *Oxytricha* suggested a transposase can recognize the DNA at the break sites with relatively relaxed sequence specificity (46). Overall, the diverse mechanisms that have evolved to fulfill the function of generating DSBs, as evidenced by the variations in both nematodes and ciliates, underscore the complexity of the PDE process.

### The SFE as a genetic tool to manipulate chromosomes and karyotypes

The insertion of the SFE in the middle of the sex chromosome (Fig. 5) demonstrated that we can use the SFE as a tool to change the genome and create new somatic karyotypes in *O. tipulae*. This is feasible because nematode chromosomes are holocentric, with functional centromeres distributed along the length of the chromosomes (37,38). As long as there are enough functional centromeres on the chromosome fragments, in theory, the fragments capped with telomeres can become independent somatic chromosomes (Fig. 5). In *Ascaris*, we previously showed that centromere deposition is dynamic during development and that centromere reorganization facilitates the elimination of DNA (62).

The mutant with a split X chromosome did not exhibit any obvious phenotype. This could be because the split occurred at a site flanking heterochromatic regions, allowing the formation of two typical nematode chromosomes without significantly disrupting genome organization or gene expression. Interestingly, the insertion site is at the boundary of two distinct ancestral linkage groups (Nigon E and X) (30), suggesting that it could be close to the site of an ancient chromosome fusion event. The two split chromosomes are nearly equal in size (5 to 6 Mbs), which are expected to contain many functional centromeres for their proper segregation during mitosis. Furthermore, nematode sex chromosomes appear to be able to tolerate and have undergone more extensive fusions and fissions compared to autosomes (26,30,40). Thus, our choice of using the sex chromosome and the positioning of the SFE insert may have contributed to our success. It would be intriguing to determine in the future whether SFE can also split autosomes and generate distinct somatic karyotypes.

The *O. tipulae* SFE system also offers a chance to develop genetic tools to manipulate other nematode chromosomes and beyond. By understanding the molecular players involved in SFE recognition, DSB generation, and DSB end repair, one could create a system to place the SFE at specific regions of the genome to introduce breaks and alter karyotypes. This approach could be applied to other nematodes with holocentric chromosomes and further developed and adapted to other eukaryotic systems with monocentric chromosomes, particularly if the changes are to be made at the chromosome ends. Ultimately, understanding the molecular mechanisms of the SFE system could lead to the development of a novel toolkit that may add to the growing array of tools for editing and engineering eukaryotic chromosomes (63–66).

## Conclusion

Our work reveals the sequence determinants of the SFE in *O. tipulae* that facilitate DNA breaks and telomere addition during PDE. We demonstrated that the SFE is both necessary and sufficient for PDE. We identified several sequence requirements of the SFE, including those for telomere priming. Our study further illustrates the remarkable flexibility in the mechanisms that have evolved to execute PDE in diverse species. Given the increasing number of organisms discovered to have PDE, we believe that understanding the mechanisms of PDE in various systems will continue to yield novel insights into the evolutionary and developmental dynamics of genomes and chromosomes.

## Materials and Methods

### Worm culture, maintenance, and staging of embryos and larvae

The *Oscheius tipulae* CEW1 strain, originally isolated by Carlos E. Winter in Brazil, was obtained from the Caenorhabditis Genetics Center (CGC, https://cgc.umn.edu/). Worms were cultured on standard Nematode Growth Medium (NGM) agar plates seeded with *Escherichia coli* OP50 at room temperature (22 °C) and were transferred to fresh plates every four days to maintain healthy populations for CRISPR injections. For staging of embryos (for END-seq and CUT&RUN), a liquid-culture approach was used to synchronize the worm population (29). Briefly, dauer were collected after the cultured worms were treated with 1% SDS followed by sucrose flotation. The dauer were then seeded on rich agar plates and incubated at room temperature for 36 to 44 hours. During this period, the worms were monitored to determine the timing of their egg-laying. Early embryos (mostly mix of 2-16 cells) were collected by bleaching the worms (67). These embryos were further developed to reach the desired embryo stages. For staging of larvae (for CUT&RUN and Hi-C), early embryos or mixed embryos were developed and arrested in Virgin S Basal (VSB) medium (100 mM NaCl, 5.7 mM K2HPO4, 44.1 mM KH2PO4). The arrested L1 larvae were seeded on NGM plates and incubated for five hours at 20°C to establish a synchronized population of L1 larvae.

### CRISPR

CRISPR-Cas9-based genome editing was adapted from our previous work (29). All CRISPR-Cas9 reagents, including crRNAs, tracrRNA, and Cas9 protein (Cat# 1081058), were obtained from Integrated DNA Technologies (IDT). Locus-specific crRNAs were selected based on combined scoring criteria using IDT’s CRISPR design tool and CRISPRscan (68), with only candidates scoring ≥40 (on a 0–100 scale) in both tools considered for further evaluation. All crRNAs, repair template sequences, and PCR primers are listed in Table S1.

Two ribonucleoprotein (RNP) complexes were prepared for injection. The co-CRISPR selection complex targeted the *O. tipulae* rol-6 gene to induce a dominant roller phenotype (69). This complex was assembled by mixing 0.5 µL of 10 mg/mL Cas9 protein with 4.5 µL of IDT duplex buffer, followed by the addition of 2.5 µL of 100 µM tracrRNA and 2.5 µL of 100 µM rol-6 crRNA. The locus-specific RNP complex targeting the SFE motif was prepared by mixing 0.5 µL of 10 mg/mL Cas9 with 3.5 µL duplex buffer, then adding 4.0 µL of 100 µM tracrRNA and 1.0 µL of 200 µM of each flanking crRNA. Both RNP complexes were incubated at 37 °C for 15 minutes to promote complex formation, after which 1.0 µL of 100 µM repair template was added to each mix to facilitate homology directed repair. The two RNP complexes were combined and centrifuged at 10,000 g for 2 minutes. Lipofectamine (Invitrogen, Cat# 13778030) was added to the supernatant at a final concentration of 3% (v/v), and the mixture was incubated at room temperature for 20 minutes before being loaded into injection needles.

Microinjections were performed using a Zeiss inverted microscope Axio Observer with DIC optics and a Narishige MMO 203 manipulator and IM400 injector, set to an input pressure of 70 psi and an injection pressure of 20 psi. Capillary needles (World Precision Instruments, Cat# 1B100F-4) were pulled using a Sutter P-1000 puller with settings at heat 490 (ramp minus 10), pull 90, velocity 80, delay 140; pressure was set to 490.

Well-fed, 4-day-old young adult *O. tipulae* were immobilized on dried 2% agarose pads and covered with halocarbon oil (Sigma, Cat# 9002839) for injection. Following microinjection, worms were gently released into M9 buffer and transferred to NGM plates seeded with *E. coli* OP50. Injected worms were incubated at 25 °C and screened after 3–5 days for F1 progeny exhibiting the dominant roller phenotype. In our experience, about 5–10% of injected P_0_ animals produced F1 roller progeny. F1 rollers were allowed to self-fertilize, and genomic DNA from pools of ten F2 progeny was screened by PCR to detect the desired CRISPR-Cas9-induced mutation (70). Putative mutants were isolated as single F2 animals, and their progeny were screened for homozygosity of the targeted edit by PCR. Homozygotes were confirmed by Sanger sequencing.

To obtain the desired mutant that alters the 1-4 nucleotides of the consensus sequence from TTTA to TTAT (Fig. 2A), we injected a total of 272 worms in four batches. We generated 17 independent roller lines and a total of 39 rollers. However, we were unable to recover any lines that harbored the desired TTAT mutation. In contrast, for most other CRISPR mutants, on average, 3-4 independent roller lines would produce the desired mutation.

### Mutant strains nomenclature

*O. tipulae* mutants were named following the *C. elegans* genetic nomenclature guidelines (71). Our laboratory allele designation “Pde” was used as a prefix, followed by either Df (for deletions or deficiencies), Dp (for duplication or insertion), or Sb (for sequence substitutions), along with the size (in base pairs) and the targeted site. For example, a 29 bp substitution at the native SFE motif with the consensus at the left arm of chromosome II is designated PdeSb29[WT-Con-2L]. An insertion of the consensus SFE inward at the left arm of chromosome II is designated PdeDp29[Con-1kb-2L]. See Table S2 for a list of mutant IDs, strain names, and a brief description of the mutants generated in this study.

### Genomic DNA extraction and genome sequencing

*O. tipulae* mutants were grown on 150 mm rich agar plates seeded with *E. coli* strain NA22. When the bacterial lawn was nearly exhausted, worms were washed off the plates using M9 buffer and pelleted by centrifugation at 170 x g for 1 min. The worm pellets were washed multiple times with M9 buffer to remove microbial contaminants (72). High molecular weight genomic DNA was extracted using Genomic-tips (Qiagen, Cat# 10223) according to the manufacturer’s protocol. DNA libraries were prepared using the NEBNext Ultra II DNA Library Prep Kit for Illumina (NEB, Cat# E7645S) and sequenced on an Illumina NovaSeq 6000 platform. The worms were a diverse mix of embryos, larvae, and adults. The sequencing results revealed that the percentage of germline DNA in the samples varied between 5% and 20%.

### END-seq

Staged embryos (8-32 cells) were used to create END-seq libraries using an adapted protocol (73,74). Briefly, embryos were embedded in agarose plugs to protect the DNA from exogenous breaks. DSBs were blunted with exonuclease VII (NEB, Cat# R0630) and exonuclease T (NEB, Cat# M0625). Blunt ends were A-tailed and capped with END-seq adaptor 1, a biotinylated hairpin adaptor. DNA was liberated from the plugs and sheared to ~200-300 bp with a Covaris M220 focused-ultrasonicator (130 μL tube, peak power 50, duty 16, cycles/burst 200 for 420 seconds). DNA fragments containing END-seq adaptor 1 were isolated with Dynabeads™ MyOne™ Streptavidin C1 (Invitrogen, Cat# 65001). The other ends broken by sonication were repaired and A-tailed with END-seq adaptor 2. Hairpins were digested with USER (NEB, Cat# M5505), and the DNA was amplified with Illumina TruSeq primers and barcodes. The libraries were sequenced with an Illumina NovaSeq 6000.

### CUT&RUN

*O. tipulae* embryos were homogenized with a metal dounce tissue grinder and larvae were ground to a fine powder using a mortar and pestle with liquid nitrogen. Each CUT&RUN reaction used ~50 μL of packed embryos or tissue powder. CUT&RUN was performed using the CUTANA ChIC-CUT&RUN kit (Epicypher Cat#14-1048) according to the manufacturer’s protocol. For each reaction, 0.5 μg of antibody against H3K9me3 (CMA318) or H3K4me3 (CMA304) was used. Library preparation was done using a 2S Plus DNA Library Kit from Swift/IDT (Swift Cat# 21024). The libraries were sequenced with an Illumina NovaSeq 6000.

### Hi-C

Synchronous L1 larvae from the wild-type strain and the mutant strain harboring the SFE consensus sequence in the middle of the X chromosome (*con-X-middle*) were used to construct two Hi-C libraries using the Arima-Hi-C 2.0 kit (Arima, Cat# A410110), as per the manufacturer’s protocol. The larvae were fixed using the low input protocol, with the following modification: larval membranes were disrupted by forcing cells through a metal dounce in 5 mL of PBS + TC fixative buffer kept chilled on ice. After cross-linking and digestion, samples were fragmented using the Covaris M220 ultrasonicator with the following settings: temperature = 7°C, peak incident power = 75 W, duty factor = 5%, cycles per burst = 200, time = 70 s. Samples were sonicated to fragments of ~500 bp, confirmed using a TapeStation 4200 (Agilent). Size selection was performed using AMPure XP magnetic beads (Beckman Coulter, Cat# A63881). Illumina libraries were constructed using the NEBNext Ultra II DNA Library Prep Kit (NEB Cat# E7645S). Libraries were amplified using the KAPA Library Amplification kit (Roche Cat# KK2620) according to the manufacturer’s protocols. The libraries were sequenced with an Illumina NovaSeq 6000.

## Data analysis

### SFE alignment and motif analysis

Twelve chromosomal break site sequences at the junctions of retained and eliminated DNA were extracted from the *O. tipulae* CEW1 reference genome (30). Multiple sequence alignment was initially performed using ClustalW algorithm in Jalview (version 2.11.4.1) (75). Manual adjustments were applied to improve alignment consistency at the conserved GGC/GCC boundaries (Fig. 1A). For the consensus, the extra guanine (G) residues in the middle of the SFE sites of chromosomes IIIR, VL, and VR were removed based on the low frequencies of 4-nt spacer among all SFEs. The sequence motif was generated using the ggseqlogo package in R (76).

### Genomic data mapping and analysis

Genomic reads from *O. tipulae* SFE mutants were aligned to the *O. tipulae* telomere-to-telomere reference genome (30) or the corresponding mutant genomes using Bowtie2 (77). The resulting SAM files were converted to BAM format and sorted using SAMtools (78). Read coverage across the genome was calculated using BEDTools to generate a BEDGRAPH file (79), which was subsequently uploaded to the UCSC Genome Browser (https://genome.ucsc.edu/s/jianbinwang/OscheiusSFE) via custom track data hubs for visualization (80).

### Telomere and END-seq analysis

END-seq reads (R1 file only) were first mapped to the *O. tipulae* reference genome (30) or the corresponding mutant genomes with Bowtie2 (77) and processed with SAMtools (78). The 5’ position of each read was mapped, and its strand was determined with genomecov in BEDTools (79). This analysis determines the DSB and its resected ends and confirms END-seq enriches DSBs and telomeres. Following that, telomere-unique reads were identified as sequences containing two or more sequential telomeric repeats (TTAGGCTTAGGC). All telomere-unique reads were converted to the G-rich strand (TTAGGC). Fastx_clipper was then used to trim the telomeric sequences. These trimmed reads were mapped to the genome, and the telomere-unique junctions were used to identify the telomere addition sites. False positive reads originating from germline genomic regions containing telomere repeats were filtered out.

### CUT&RUN, ATAC-seq, and RNA-seq data analysis

Chromatin profiling datasets, including CUT&RUN for H3K9me3, H3K4me3, ATAC-seq, and RNA-seq, were processed using a uniform alignment and peak calling process. For alignment, reads were first mapped to the reference genome using Bowtie2 (77). The resulting SAM files were converted to BAM format and sorted using SAMtools (78). To reduce PCR amplification artifacts, duplicate reads were removed using Picard’s MarkDuplicates tool http://broadinstitute.github.io/picard/. For normalization across chromatin marks for different staged embryos, duplicate-free BAM files were down sampled to a fixed depth of 10 million uniquely mapped reads using SAMtools with a fixed random seed to ensure reproducibility.

Peak calling was performed on normalized BAM files using MACS3 (81). Peaks were identified using a stringent q-value cutoff of 0.01, and both narrowPeak and summits.bed files were retained for each dataset. To assess chromatin feature enrichment within specific genomic contexts, peaks were intersected with a curated set of 31 predefined genomic regions (flanking 500 bp at both sides of the sequence), which included canonical SFE sites, alternative SFE sites, FIMO-predicted sequences, and regions used for SFE insertions in the retained region (see Fig. 6C-D). Intersections were performed using BEDTools (79) to determine if the SFE regions are overlapping with the defined peaks. For control, we used 31 randomly generated regions across the genome using the *random* command in BEDTools (79).

### Visualization of telomere addition using Integrative Genomics Viewer (IGV)

Raw genomic reads from the mutant strains were processed using a standard data processing procedure for quality control and alignment to the *O. tipulae* reference genome (30) or the corresponding mutant genomes. For IGV viewing purpose, adapter trimming and quality filtering were performed using Trimmomatic in paired-end mode to remove adapter contamination with the following parameters: ILLUMINACLIP:TruSeq3-PE-2.fa:2:30:10:1:True and MINLEN:15. Trimmed reads were assessed using FastQC (82) to evaluate post-trimming quality, sequence content, and duplication levels. Only the trimmed and paired reads were used for downstream analysis (83).

They were aligned to the mutant reference genome using Bowtie2 (77) with the --local and -X 5000 parameters, allowing for soft clipping and accommodating larger insert sizes, respectively. The resulting SAM files were converted to BAM format using SAMtools (78), then sorted by genomic coordinate and indexed to produce IGV-compatible .bam and .bai files. These BAM files were visualized using the IGV (84) to inspect read coverage, telomere additions, and evaluate the outcomes of genome changes in the SFE mutants.

### Hi-C analysis

Paired-end Hi-C reads were aligned to the *Oscheius* reference genome using Bowtie2 (77), and contact matrices were generated with HiC-Pro (85,86). AllValidPairs files were converted to .hic format using hicpro2juicebox from HiC-Pro for downstream analysis. Standard HiC-Pro quality control metrics were computed for all samples, including mapping rates, valid interaction percentages, and trans/cis interaction ratios. Contact heatmaps were generated at a resolution of 10 kb using custom python codes and ICE-normalized data. Datasets were normalized between each other using counts per million reads (cpm).

## Supplementary materials

### List of supplemental tables

Table S1. List of mutants generated in this study.

Table S2. List of sequences for crRNAs, repair templates, and PCR primers.

### List of supplemental figures

Figure S1. IGV view of genomic reads for the *WT-Con* and *Con-Swap* mutants.

Figure S2. IGV view of genomic reads for the mutants with 4-bp mutated sequences.

Figure S3. IGV view of genomic reads for the mutants at the telomere addition site.

Figure S4. IGV view of genomic reads for the mutants at 1kb and 10kb adjacent to native SFE at chrII-L

## Data availability

All sequencing data are deposited in the NCBI SRA database with accession number PRJNA1425020. The CUT&RUN, END-seq, Hi-C, and genome sequencing data are also available in the NCBI GEO database with accession numbers GSE323393, GSE323395, GSE323396, and GSE323397, respectively. The data are also accessible in UCSC Genome Browser track data hubs with this link: https://genome.ucsc.edu/s/jianbinwang/OscheiusSFE.

## Acknowledgment

We thank CGC, funded by NIH Office of Research Infrastructure Programs (P40 OD010440), for the *O. tipulae* strain. We thank Jan Hudson and undergraduate students Daniella Morales, Connie Bailey, Bobbie Moore, Matthew Hughes, and Jessica Garner for helping with plates and worm maintenance; Nathan Fawver for assistant in CUT&RUN; and the University of Colorado Anschutz Medical Campus Genomics Core for sequencing services. We also thank the members of the Wang lab for their discussions and Albrecht von Arnim, Larry McReynolds, and Dick Davis for comments and critical reading of the manuscript.

## Author contributions

T.C.D. and J.W. conceived the study; J.S. and T.C.D. did worm culture and maintenance with the help of M.I., V.T., M.A., A.W., and H.L.; M.A., V.T., and T.C.D. performed CRISPR injection with the help of M.I., A.W., and H.L.; J.S. and T.C.D. screened for homozygosity and established mutant lines; J.S. and T.C.D. sequenced the mutant genomes; B.E. carried out END-seq; R.O. did CUT&RUN; J.R.S. performed Hi-C; J.S. and J.W. carried out data analysis with the help of B.E. and J.R.S.; J.W. wrote the initial manuscript draft; J.S., T.C.D., and J.W. edited the manuscript; and J.W. provided supervision, project administration, and funding.

