## Supplemental Files for "A sequence motif for DNA double-strand break and telomere healing during programmed DNA elimination"

**Table S1. List of mutants generated in this study**

| Mutant ID | Mutant strain name | Brief description of the mutant | PDE Efficiency |
|---|---|---|---|
| WT-con | *PdeSb29[WT-Con-2L]* | Replacement of the native SFE at chrII-L with the SFE consensus (Fig. 1D) | 100% |
| Con-Swap | *PdeSb29[Con-Swap-2L]* | Replacement of the native SFE at chrII-L with the swapped SFE consensus (Fig. 1F) | 100% |
| AGGT>GATG | *PdeSb29[Con-4-7-GATG-2L]* | A mutated motif at nucleotides 4–7 of the SFE consensus (Fig. 2B) | 30% |
| TTTG>TGTT | *PdeSb29[Con-7-10-TGTT-2L]* | A mutated motif at nucleotides 7–10 of the SFE consensus (Fig. 2B) | 100% |
| GGCT>CGTG | *PdeSb29[Con-11-14-CGTG-2L]* | A mutated motif at nucleotides 11–14 of the SFE consensus (Fig. 2B) | 0% |
| CTTA>TCAT | *PdeSb29[Con-13-16-TCAT-2L]* | A mutated motif at nucleotides 13–16 of the SFE consensus (Fig. 2B) | 0% |
| GCC>CCC | *PdeSb29[Con-17-19-CCC-2L]* | A mutated motif at nucleotides 17–19 of the SFE consensus (Fig. 3B) | 11% |
| GCC>GGC | *PdeSb29[Con-17-19-GGC-2L]* | A mutated motif at nucleotides 17–19 of the SFE consensus (Fig. 3B) | 75% |
| Con-1kb | *PdeDp29[Con-1kb-2L]* | Insertion of consensus at 1-kb towards the retained side of the native ChrII-L SFE (Fig. 4) | 100% |
| Con-10kb | *PdeDp29[Con-10kb-2L]* | Insertion of consensus at 10-kb towards the retained side of the native ChrII-L SFE (Fig. 4) | 100% |
| Con-X-middle | *PdeDp29[Con-X-middle]* | Mutant that harbors the SFE consensus in the middle of the X chromosome (Fig. 5A) | 100% |

### Table S2. List of sequences for crRNAs, repair templates, and PCR primers

**Common Crispr Reagents**

| Mutant Name | ID (Name) | Sequence |
|---|---|---|
| ***Roller*** | rol-6-crRNA | CGUGUCCGUCGUCAGUAUGG |
| ***rol-6*** | rol-6-repair-template | GGGAGATATGGTCAAGCTGGGAGCTGGATCTGCCTCAAACCGTGTCAGATGCCAGTA<br>TGGAGGATACGGTGCTAGTGGAGTTCAGCCTCAGCCTCCTCAG |
| | crRNA-2L_Fwd | UAGAAGGCGUUUAUAA |
| | crRNA-2L_Rev | UGCCCAUAGGCUUUUGU |

**Mutants Specific Reagents**

| Mutant Name | ID (Name) | Sequence | PCR Product size |
|---|---|---|---|
| ***PdeSb29[WT-Con-2L]*** | WT-cons Repair-template | CAACGAGGTCTCCAGCTGAAGAAATTATTACAATAGATGTCTTTGTTTTAGAAGGCTG<br>TTTATAACTTTGACTTTAGGTTTGGGCTTAGCCAAAGCCCAAAATATTGTGTTGTAATTT<br>ATCACGTGTGCCCATAGGCTTTTGTGACACACCACCCTCAAAAATTTTAGCTTTCTATC<br>CAAGTGCCATTGAC | |
| | WT-cons Fwd-primer | AGGTTTGGGCTTAGCCAAAGCCC | |
| | WT-cons Rev-primer | GGAGTCAATGGCACTTGGATAG | 119bp |
| ***PdeSb29[Con-1-4-TTAT-2L]*** | TTTA-TTAT Repair-template | CAACGAGGTCTCCAGCTGAAGAAATTATTACAATAGATGTCTTTGTTTTAGAAGGCTG<br>TTTATAACTTTGACTTATGGTTTGGGCTTAGCCAAAGCCCAAAATATTGTGTTGTAATTT<br>ATCACGTGTGCCCATAGGCTTTTGTGACACACCACCCTCAAAAATTTTAGCTTTCTATC<br>CAAGTGCCATTGAC | |
| | TTTA-TTAT Fwd-primer | AGGTTTGGGCTTAGCCAAAGCCC | |
| | TTTA-TTAT Rev-primer | GGAGTCAATGGCACTTGGATAG | 119bp |
| ***PdeSb29[Con-4-7-GATG-2L]*** | AGGT-GATG Repair-template | CAACGAGGTCTCCAGCTGAAGAAATTATTACAATAGATGTCTTTGTTTTAGAAGGCTG<br>TTTATAACTTTGACTTTGATGTTGGGCTTAGCCAAAGCCCAAAATATTGTGTTGTAATTT<br>ATCACGTGTGCCCATAGGCTTTTGTGACACACCACCCTCAAAAATTTTAGCTTTCTATC<br>CAAGTGCCATTGAC | |
| | AGGT-GATG Fwd-primer | GATGTTGGGCTTAGCCAAAGCCC | |
| | AGGT-GATG Rev-primer | GGAGTCAATGGCACTTGGATAG | 119bp |
| ***PdeSb29[Con-7-10-TGTT-2L]*** | TTTG-TGTT Repair-template | CAACGAGGTCTCCAGCTGAAGAAATTATTACAATAGATGTCTTTGTTTTAGAAGGCTG<br>TTTATAACTTTGACTTTAGGTGTTGGCTTAGCCAAAGCCCAAAATATTGTGTTGTAATTT<br>ATCACGTGTGCCCATAGGCTTTTGTGACACACCACCCTCAAAAATTTTAGCTTTCTATC<br>CAAGTGCCATTGAC | |
| | TTTG-TGTT Fwd-primer | AGGTGTTGGCTTAGCCAAAGCCC | |
| | TTTG-TGTT Rev-primer | GGAGTCAATGGCACTTGGATAG | 119bp |
| ***PdeSb29[Con-11-14-CGTG-2L]*** | GGCT_CGTG Repair-template | CAACGAGGTCTCCAGCTGAAGAAATTATTACAATAGATGTCTTTGTTTTAGAAGGCTG<br>TTTATAACTTTGACTTTAGGTTTGCGTGTAGCCAAAGCCCAAAATATTGTGTTGTAATTT<br>ATCACGTGTGCCCATAGGCTTTTGTGACACACCACCCTCAAAAATTTTAGCTTTCTATC<br>CAAGTGCCATTGAC | |
| | GGCT_CGTG Fwd-primer | AGGTTTGCGTGTAGCCAAAGCCC | |
| | GGCT_CGTG Rev-primer | GGAGTCAATGGCACTTGGATAG | 119bp |
| ***PdeSb29[Con-13-16-TCAT-2L]*** | CTTA-TCAT Repair-template | CAACGAGGTCTCCAGCTGAAGAAATTATTACAATAGATGTCTTTGTTTTAGAAGGCTG<br>TTTATAACTTTGACTTTAGGTTTGGGTCATGCCAAAGCCCAAAATATTGTGTTGTAATTT<br>ATCACGTGTGCCCATAGGCTTTTGTGACACACCACCCTCAAAAATTTTAGCTTTCTATC<br>CAAGTGCCATTGAC | |
| | CTTA-TCAT Fwd-primer | AGGTTTGGGTCATGCCAAAGCCC | |
| | CTTA-TCAT Rev-primer | GGAGTCAATGGCACTTGGATAG | 119bp |
| ***PdeSb29[Con-17-19-CCC-2L]*** | GCC-CCC Repair-template | CAACGAGGTCTCCAGCTGAAGAAATTATTACAATAGATGTCTTTGTTTTAGAAGGCTG<br>TTTATAACTTTGACTTTAGGTTTGGGCTTACCCAAAGCCCAAAATATTGTGTTGTAATTT<br>ATCACGTGTGCCCATAGGCTTTTGTGACACACCACCCTCAAAAATTTTAGCTTTCTATC<br>CAAGTGCCATTGAC | |
| | GCC-CCC Fwd-primer | AGGTTTGGGCTTACCCAAAGCCC | |
| | GCC-CCC Rev-primer | GGAGTCAATGGCACTTGGATAG | 119bp |
| ***PdeSb29[Con-17-19-GGC-2L]*** | GCC-GGC Repair-template | CAACGAGGTCTCCAGCTGAAGAAATTATTACAATAGATGTCTTTGTTTTAGAAGGCTG<br>TTTATAACTTTGACTTTAGGTTTGGGCTTAGGCAAAGCCCAAAATATTGTGTTGTAATT<br>TATCACGTGTGCCCATAGGCTTTTGTGACACACCACCCTCAAAAATTTTAGCTTTCTAT<br>CCAAGTGCCATTGAC | |
| | GCC-GGC Fwd-primer | AGGTTTGGGCTTAGGCAAAGCCC | |
| | GCC-GGC Rev-primer | GGAGTCAATGGCACTTGGATAG | 119bp |
| ***PdeSb29[Con-Swap-2L]*** | Cons-Swap Repair-template | CAACGAGGTCTCCAGCTGAAGAAATTATTACAATAGATGTCTTTGTTTTAGAAGGCTG<br>TTTATAACTTTGACTTTGGGCTTTGGCTAAGCCCAAACCTAAAATATTGTGTTGTAATTT<br>ATCACGTGCATAGGCTTTTGTGACACACCACCCTCAAAAATTTTAGCTTTCTATCCAAG<br>TGCCATTGAC | |
| | Cons-Swap Fwd-primer | CCTCCGCTGAGCGTCTTATT | |
| | Cons-Swap Rev-primer | CACAAAAGCCTATGCACGTG | 212bp |
| ***PdeDp29[Con-inward-1kb-2L]*** | crRNA-Con-inward-1kb | UUGGAGGCGUUUCUCGUUUU | |
| | Con-inward-1kb Repair-template | CTTTGTGTAACTTTGTATCTCTGACTCTGTCTCTATCTTTCTCTTCTGATTCTATATTTGC<br>CCTCCAAAAACACAATATTTTGGGCTTTGGCTAAGCCCAAACCTAAAGTCAAAGTTATA<br>AACGAGAAACGCCTCCAATTTTCGGCTTTACGCCCGCTTTATCATCATTCCCTCCTGGT<br>ATTAAGGACGTCTC | |
| | Con-inward-1kb Fwd-primer | TGCGACTGAAATGGCTTTTTGAA | |
| | Con-inward-1kb Rev-primer | TGATAAAGCGGGCGTAAAGC | 310bp |
| ***PdeDp29[Con-inward-10kb-2L]*** | crRNA-Con-inward-10kb | GCAAGAUCAAUGAGGAAGAA | |
| | Con-inward-10kb Repair-template | GACTTTTTCTCGCGTACCAACTGATCCCTTCTCTTCCATTTTTCTGAGCTCTGCCCTTCA<br>TAAATTACAACACAATATTTTGGGCTTTGGCTAAGCCCAAACCTAAAGTCAAAGTTATA<br>AACAGCCTTCTTTCCTCATTGATCTTGCTATTTCTGATCCCTTTTTTGCATTTTATTGAAG<br>GATAACACCAAATGCATAC | |
| | Con-inward-10kb Fwd-primer | TTTAGGCGCGACTTTTTCTCG | |
| | Con-inward-10kb Rev-primer | CAGCAAACAAATCATGGTGGT | 315bp |
| ***PdeDp29[Con-X-middle]*** | crRNA-Con-X-middle | CUGGAUUUGCACUUCUCCAC | |
| | Con-X-middle Repair-template | TGAACTCCACACGTACGACAGTTCACAGTGGCAGCGGCTATTTTTAAGTAGGATTGTG<br>TCCGGTGATAAATTACAACACAATATTTTGGGCTTTGGCTAAGCCCAAACCTAAAGTC<br>AAAGTTATAAACAGCCTTCTGAGAAGTGCAAATCCAGGCATTATATTTCAGCCCGAGA<br>GACCAGTGATGGACTAAAGAATTCAC | |
| | Con-X-middle Fwd-primer | TGCCCTTTGCGATATCCTCG | |
| | Con-X-middle Rev-primer | CGAGCAACGATGATGGCTTC | 326bp |

RNA sequences are color in blue

Figure S1

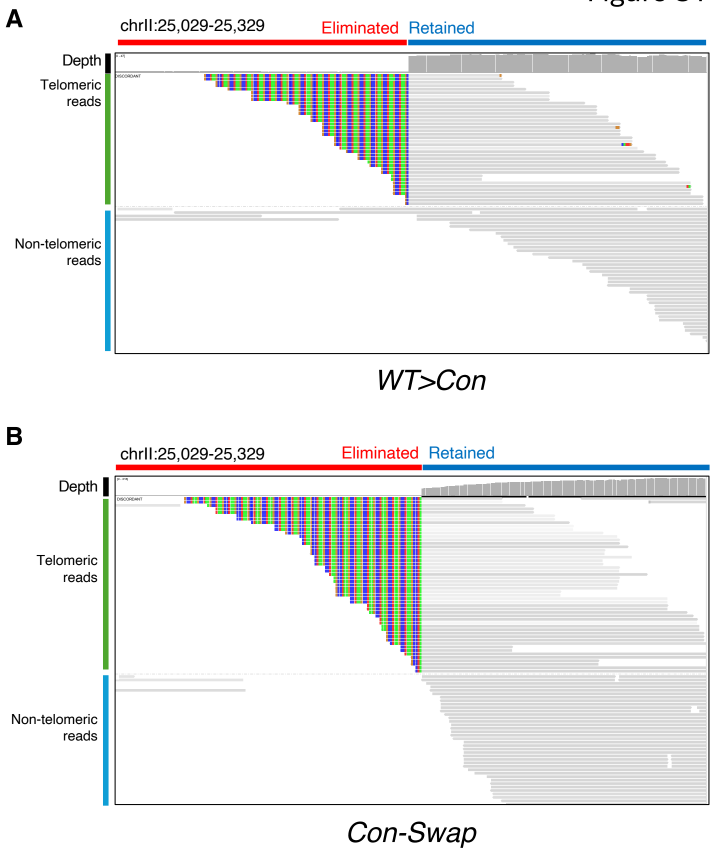


**Figure S1. IGV view of genomic reads for the *WT-Con* and *Con-Swap* mutants.** The horizontal red bar at the top marks the eliminated region, while the blue bar indicates the retained region. Reads with new telomere addition (TTAGGCs) are highlighted with colored bases and marked with a green vertical line. Non-telomeric (unchanged germline) reads are marked with a blue vertical line.

Figure S2

A

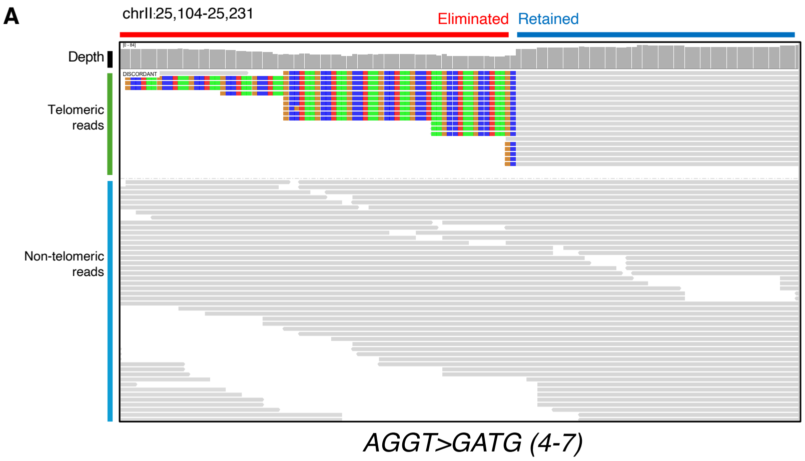


B

| Chr | Germ | Soma | Total | %Germline |
|---|---|---|---|---|
| chrIL | 2 | 48 | 50 | 4 |
| chrIR | 2 | 56 | 58 | 3.4 |
| chrIIL | 47 | 19 | 66 | 71.2 |
| chrIIR | 3 | 40 | 43 | 7.0 |
| chrIIIL | 1 | 48 | 49 | 2.0 |
| chrIIIR | 1 | 56 | 57 | 1.8 |
| chrIVL | 10 | 37 | 47 | 21.3 |
| chrIVR | 1 | 56 | 57 | 1.8 |
| chrVL | 0 | 37 | 37 | 0 |
| chrVR | 6 | 67 | 73 | 8.2 |
| chrXL | 2 | 45 | 47 | 4.3 |
| chrXR | 1 | 50 | 51 | 2.0 |
| Total count (excluding chrIIL) | 29 | 540 | 569 | 5.1 |

C

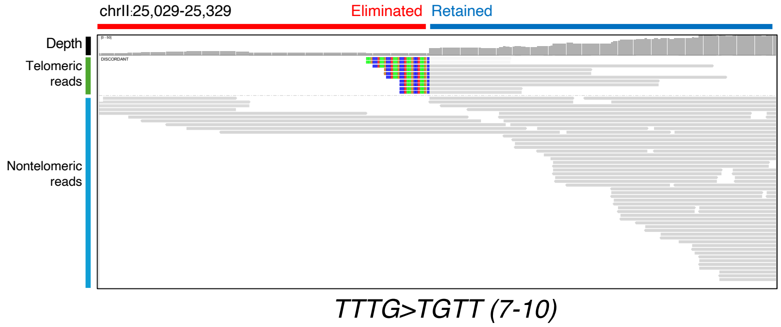


D

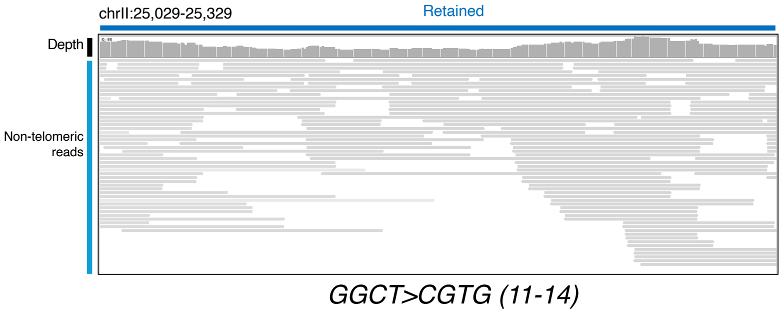


E

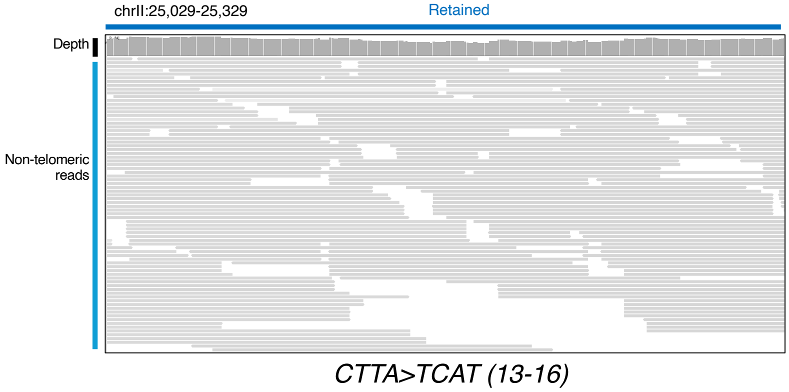


**Figure S2. IGV view of genomic reads for the mutants with 4-bp mutated sequences.** The legend is the same as in Figure S1. The read counts for the 12 canonical sites in the mutant strain AGGT-GATG (4-7) that exhibit 30% partial PDE efficiency are listed in **B**.

Figure S3

A

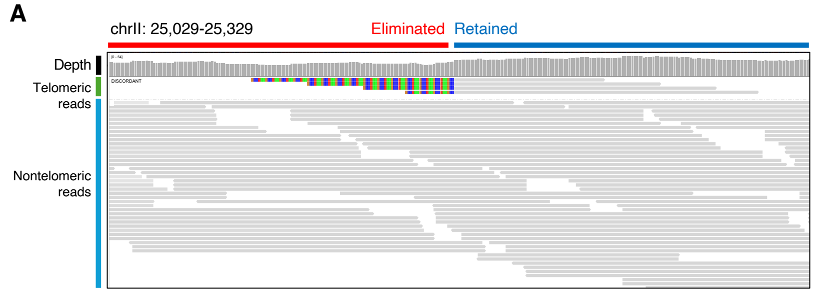


*GCC>CCC (17-19)*

B

| Chr | Germ | Soma | Total | %Germline |
|---|---|---|---|---|
| chrIL | 3 | 28 | 31 | 9.7 |
| chrIR | 2 | 37 | 39 | 5.1 |
| chrIIL | 37 | 4 | 41 | 90.2 |
| chrIIR | 3 | 36 | 39 | 7.7 |
| chrIIIL | 3 | 32 | 35 | 8.6 |
| chrIIIR | 5 | 32 | 37 | 13.5 |
| chrIVL | 7 | 28 | 35 | 20.0 |
| chrIVR | 2 | 33 | 35 | 5.7 |
| chrVL | 0 | 20 | 20 | 0 |
| chrVR | 3 | 45 | 48 | 6.3 |
| chrXL | 5 | 30 | 35 | 14.3 |
| chrXR | 1 | 25 | 26 | 3.8 |
| Total count (excluding chrIIL) | 34 | 346 | 380 | 9.0 |

C

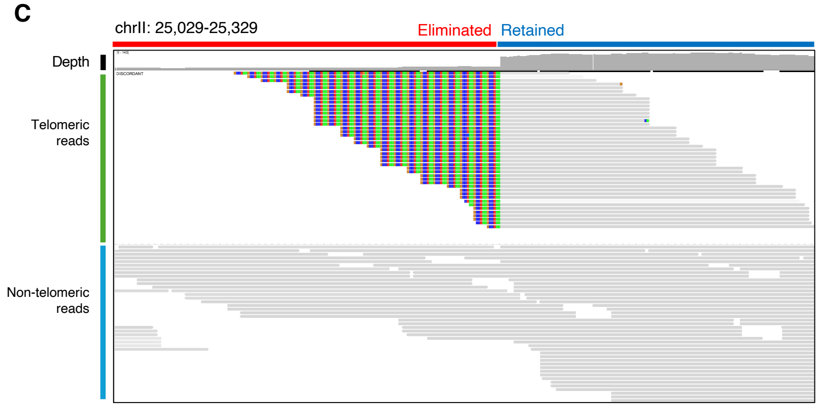


*GCC>GGC (17-19)*

D

| Chr | Germ | Soma | Total | %Germline |
|---|---|---|---|---|
| chrIL | 12 | 49 | 61 | 19.7 |
| chrIR | 13 | 101 | 114 | 11.4 |
| chrIIL | 21 | 43 | 64 | 32.8 |
| chrIIR | 2 | 58 | 60 | 3.3 |
| chrIIIL | 2 | 55 | 57 | 3.5 |
| chrIIIR | 9 | 77 | 86 | 10.5 |
| chrIVL | 9 | 50 | 59 | 15.3 |
| chrIVR | 9 | 61 | 70 | 12.9 |
| chrVL | 1 | 53 | 54 | 1.9 |
| chrVR | 12 | 102 | 114 | 10.5 |
| chrXL | 7 | 36 | 43 | 16.3 |
| chrXR | 6 | 73 | 79 | 7.6 |
| Total count (excluding chrIIL) | 82 | 715 | 797 | 10.3 |

**Figure S3. IGV view of genomic reads for the mutants at the telomere addition site.** The legend is the same as in Figure S1. The read counts for the 12 canonical sites in the mutant strain CCC, which exhibits 11% partial PDE efficiency, and the mutant strain GGC, which exhibits 75% partial PDE efficiency, are also listed.

Figure S4

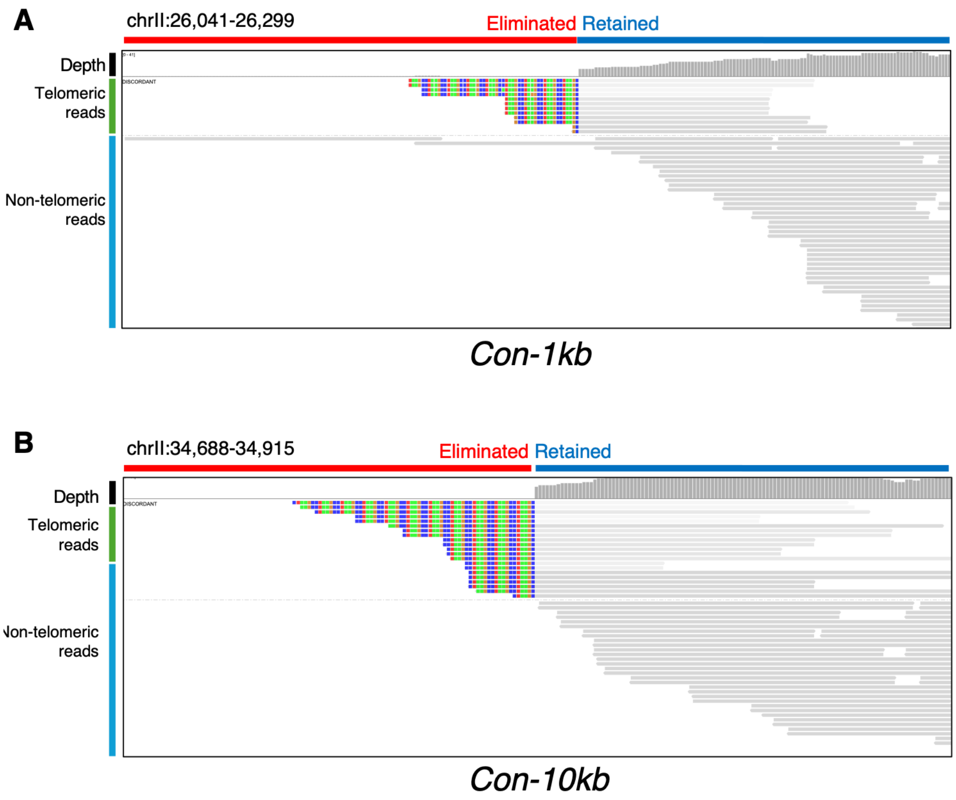


**Figure S4. IGV view of genomic reads for the mutants at 1kb and 10kb adjacent to native SFE at chrII-L.** Genomic reads for the Con-1kb and Con-10kb mutants. The legend is the same as in Figure S1.
